# BioJupies: Automated Generation of Interactive Notebooks for RNA-seq Data Analysis in the Cloud

**DOI:** 10.1101/352476

**Authors:** Denis Torre, Alexander Lachmann, Avi Ma’ayan

## Abstract

Interactive notebooks can make bioinformatics data analyses more transparent, accessible and reusable. However, creating notebooks requires computer programming expertise. Here we introduce BioJupies, a web server that enables automated creation, storage, and deployment of Jupyter Notebooks containing RNA-seq data analyses. Through an intuitive interface, novice users can rapidly generate tailored reports to analyze and visualize their own raw sequencing files, their gene expression tables, or fetch data from >5,500 published studies containing >250,000 preprocessed RNA-seq samples. Generated notebooks have executable code of the entire pipeline, rich narrative text, interactive data visualizations, and differential expression and enrichment analyses. The notebooks are permanently stored in the cloud and made available online through a persistent URL. The notebooks are downloadable, customizable, and can run within a Docker container. By providing an intuitive user interface for notebook generation for RNA-seq data analysis, starting from the raw reads, all the way to a complete interactive and reproducible report, BioJupies is a useful resource for experimental and computational biologists. BioJupies is freely available as a web-based application from: http://biojupies.cloud and as a Chrome extension from the <u>Chrome Web Store</u>.

## Introduction

RNA-sequencing^1^ is a widely applied experimental method to study the biological molecular mechanisms of cells and tissues in human and model organisms. Currently, experimental biologists that perform RNA-seq experiments are experiencing a bottleneck. The raw read FASTQ files, which are relatively large (>1 GB), need to be first aligned to the reference genome before they can be further analyzed and visualized to gain biological insights. The alignment step is challenging because it is computationally demanding, typically requiring specialized hardware and software. Recently, we have developed a cost effective cloud-based alignment pipeline that enabled us to align >250,000 publicly available RNA-seq samples from the Sequence Read Archive (SRA)^2^. Here, we describe a service that enables users to upload and align their own FASTQ files, and then obtain an interactive report with complete analysis of their data delivered as a Jupyter Notebook^3^.

Inspired by the paradigm of literate programming^4^, data analysis interactive notebook environments such as Jupyter Notebooks, R-markdown^5^, knitr^6^, Observable (https://beta.observablehq.com), or Zeppelin (https://zeppelin.apache.org) have been rapidly gaining traction in computational biomedical research and other data intensive scientific fields. The concept of an interactive notebook is not new. Computing software platforms such as MATLAB and Mathematica deployed notebook style analysis pipelines for many years. However, the availability of open-source and free interactive notebooks that can run and execute in any browser make these new notebook technologies transformative.

Interactive notebooks enable the generation of executable documents that contain source code, data analyses and visualizations, and rich narrative markup text. By combining all the necessary information to rerun data analysis pipelines, and by producing reports that enable rapid interpretation of experimental results, interactive notebooks can be considered a new form of publication. Hence, interactive notebooks can transform how experimental results are exchanged in biomedical research. In this direction, several academic journals now support the Jupyter Notebook as a legitimate component of a publication, for example, the journal F1000 Research^9^, or as an acceptable format type to submit articles, for example, the journal Data Science^10^. However, generating interactive notebooks requires high level of computer programming expertise which is uncommon among experimental biologists.

The analysis of RNA-sequencing data, and the processing of large datasets produced by other omics technologies, typically requires the chaining of several bioinformatics tools into a computational pipeline. In the pipeline, the output of one tool serves as the input to the next tool. In order to enable experimental biologists to execute bioinformatics pipelines, including those developed to process and analyze RNA-seq data, several platforms have been developed, for example, GenomeSpace^11^, Galaxy^12^, and GenePattern^13^. These software platforms provide access to workflows that can run on scalable computing resources through a web interface. Hence, users with limited computational expertise can launch computationally intensive data analysis jobs with these platforms. Galaxy and GenePattern have recently integrated Jupyter^14, 15^. In addition to these platforms, several interactive web applications for analysis of RNA-seq data have been developed. Most of these platforms are implemented with the the R Shiny toolkit^16^. Instead of relying on the integration of external computational tools, these R Shiny applications analyze uploaded data by executing R code on the server side, and then displaying the results directly in the browser. Examples of such applications include iDEP^17^, START^18^, VisRseq^19^, ASAP^20^, DEApp^21^, and IRIS^22^. However, none of these tools currently allow for the automated generation of reusable and publishable reports that contain research narratives in a single interactive notebook. In addition, these platforms do not provide users with the ability to upload the raw sequencing files for processing in the cloud.

BioJupies is a web-based server application that automatically generates customized Jupyter Notebooks for analysis of RNA-seq data. BioJupies allows users to rapidly generate tailored, reusable reports from their raw or processed sequencing data, as well as fetch RNA-seq data from >5,500 studies that contain >250,000 RNA-seq samples published in the Gene Expression Omnibus (GEO)^23^. To enable such fetching, we preprocessed all the human and mouse samples profiled by the major sequencing platforms by aligning them to the reference genome via the ARCHS4 cloud computing architecture^2^. Most importantly, BioJupies enables users to upload their own FASTQ files for alignment and downstream analysis in the cloud. BioJupies is freely available from http://biojupies.cloud. BioJupies is open source on GitHub at https://github.com/MaayanLab/biojupies available for forking and contributing plug-ins to enhance the system.

## Methods

### The BioJupies interactive website

The BioJupies website is hosted on a web server running in a Docker^24^ container as a Ubuntu image that is pulled from Docker Hub. It is served using a Python-Flask framework with an nginx HTTP server, uWSGI, and a MySQL database. The front-end of the website is built using HTML5 Bootstrap, CSS3, and JavaScript. When a user requests the generation of a notebook, the website builds a JSON-formatted ‘notebook configuration’ string that contains information about the selected datasets and plug-ins. The website first queries the MySQL database to find whether a notebook with the same configuration was previously generated. If a notebook is found, the link to it is displayed on the final results page. Otherwise, the notebook configuration JSON is sent to the notebook generator server through an HTTP POST request. The server uses this information to generate a notebook. Once the notebook has been generated, the server returns a notebook link to the client for display.

### Programmatic generation of Jupyter Notebooks

The Jupyter Notebook generator server uses the information in the JSON-formatted ‘notebook configuration’ string to generate a Jupyter Notebook with the Python library *nbformat*. The server executes the notebook using the Python library *nbconvert* and then saves the notebook in a file encoded into the *ipynb* format. The file is subsequently uploaded to a Google Cloud storage bucket using the *google-cloud* Python library. The statically rendered notebook is made available on a persistent and public URL using Jupyter *nbviewer*. The Docker container that encapsulates the running server contains a copy of the BioJupies plug-in library. The BioJupies plug-in library contains all the scripts necessary to download, normalize and analyze the RNA-seq data.

### The BioJupies plug-in library

The BioJupies plug-in library is a modular set of Python and R scripts which are used to download, normalize, and analyze RNA-seq datasets for notebook generation using BioJupies. The library consists of a set of core scripts responsible for loading the RNA-seq data into the *pandas DataFrames*^25^, scripts for normalizing the data, and scripts for performing the differential gene expression analysis, currently implemented with *limma*^26^ or the Characteristic Direction^27^ methods. In addition to these, the library contains a broad range of data analysis tools organized in a plug-in architecture. Each plug-in can be used to analyze the data, and to embed interactive visualizations of the results in the output Jupyter Notebook report. Currently, the library contains 14 plug-ins divided into 4 categories (Table 1). The plug-in library is expected to be updated regularly and grow as we incorporate more computational tools submitted by developers who wish to have their tools integrated within the BioJupies framework. The BioJupies plug-in library is openly available from GitHub at: https://github.com/MaayanLab/biojupies-plugins.

**Table 1.** List of the RNA-seq data analysis plug-ins available within BioJupies. The initial BioJupies toolbox includes 14 plug-ins for the analysis of RNA-seq data, divided into four categories.

| Tool Name | Description | References |
| --- | --- | --- |
| PCA | Linear dimensionality reduction technique to visualize similarity between samples | 38, 39 |
| Clustergrammer | Interactive hierarchically clustered heatmap | 40 |
| Library Size Analysis | Analysis of readcount distribution for the samples within the dataset | 39 |
| Differential Expression Table | Differential expression analysis between two groups of samples | 26, 27 |
| Volcano Plot | Plot the logFC and logP values resulting from a differential expression analysis | 26, 27, 39 |
| MA Plot | Plot the logFC and average expression values resulting from a differential expression analysis | 26, 27, 39 |
| Enrichr Links | Links to enrichment analysis results of the differentially expressed genes via Enrichr | 41 |
| Gene Ontology Enrichment Analysis | Gene Ontology terms enriched in the differentially expressed genes (via Enrichr) | 41, 42, 39 |
| Pathway Enrichment Analysis | Biological pathways enriched in the differentially expressed genes (via Enrichr) | 41, 43, 44, 45, 39 |
| Transcription Factor Enrichment Analysis | Transcription factors whose targets are enriched in the differentially expressed genes (via Enrichr) | 41, 46, 2 |
| Kinase Enrichment Analysis | Protein kinases whose substrates are enriched in the differentially expressed genes (via Enrichr) | 41, 47, 2 |
| miRNA Enrichment Analysis | miRNAs whose targets are enriched in the differentially expressed genes (via Enrichr) | 41, 48, 49 |
| L1000CDS2 Query | Small molecules which mimic or reverse a given differential gene expression signature | 50, 39 |
| L1000FWD Query | Small molecules which mimic or reverse a given differential gene expression signature | 51 |

### Processed RNA-seq datasets

The pre-processed RNA-seq datasets available for analysis from BioJupies are directly extracted from the ARCHS4 database^2^. First, gene-level count matrices stored in the h5 format were downloaded for mouse and human from the ARCHS4 web-site (human_matrix.h5 v2 and mouse_matrix.h5 v2). Gene counts and metadata were subsequently extracted using the Python *h5py* library and packaged into 5,780 individual HDF5 files, one for each GEO Series (GSE) and GEO Platform (GPL) pair. Data packages were subsequently uploaded to a Google Cloud storage bucket using the *google-cloud* library and made available for download through a public URL.

### Processing of user-submitted RNA-seq data

Processed RNA-seq datasets are submitted from the upload page using the Dropzone JavaScript library. Uploaded files are combined with sample metadata into an HDF5 data package^28^ using the *h5py* Python library and subsequently uploaded to a Google Cloud storage bucket with the *google-cloud* library. Raw sequencing files are submitted directly to an Amazon Web Services (AWS) S3 cloud storage bucket from the raw sequencing data upload page. Once the data is uploaded, gene expression is quantified using the ARCHS4 RNA-seq processing pipeline^2^, which runs in parallel on the AWS cloud. The core component of the processing pipeline is the alignment of the raw mRNA reads to a reference genome. This process is encapsulated in deployable Docker containers that performs the alignment step with Kallisto^29^. Once gene counts have been quantified, the data is combined with sample metadata into an HDF5 package similarly to the way the processed datasets are made accessible to BioJupies for further analysis. The system allocates resources dynamically based on need to optimize cost. The use of the efficient aligner Kallisto with the optimized ARCHS4 RNA-seq processing pipeline keeps the cost negligible and the service scalable.

### The BioJupies Chrome Extension

The BioJupies Chrome extension is freely available from the Chrome Web Store at: https://chrome.google.com/webstore/detail/biojupies-generator/picalhhlpcihonibabfigihelpmpadel?hl=en-US. The Chrome extension enhances the search results pages of the GEO website at: https://www.ncbi.nlm.nih.gov/geo. When a user is visiting a search results page on GEO, the BioJupies Chrome extension extracts the accession IDs of the query returned GEO datasets. The extension then queries the BioJupies MySQL database to identify which search-returned datasets have been processed by ARCHS4^2^. For those datasets that have been processed by ARCHS4, the BioJupies Chrome extension embeds BioJupies buttons near the matching entries. When these buttons are clicked by the user, a user interface is deployed. The user interface leads the user through steps to generate the ‘notebook configuration’ JSON string. Once all information is gathered, the ‘notebook configuration’ JSON string is sent to the notebook generation server through an HTTP POST request to generate a Jupyter Notebook. Once the notebook is ready, a link to it is displayed by the Chrome extension.

## Results

### The BioJupies website

The BioJupies website enables users to generate customized, interactive Jupyter Notebooks containing analyses of RNA-seq data through an intuitive user interface. The notebooks contain executable Python code, interactive visualizations, and rich annotations that provide detailed explanations of the results. The notebooks also provide details and references about the methods used to perform the analyses. The notebooks are made available to users through a persistent sharable URL. The automatically generated notebooks can be downloaded and modified on the user’s local computer through the execution of a Docker image that contains all the data, tools and source code needed to rerun analyses.

The process of notebook generation consists of three steps. First, the user selects the data they wish to analyze through a web interface. Users can either upload their own raw or processed RNA-seq data (Supplementary Video S1), or select from over 5,500 ready-to-analyze datasets published in GEO and processed by ARCHS4. Next, the user selects from an array of computational plug-ins to analyze the data (Supplementary Video S2). Finally, the user can customize the notebook by modifying optional parameters, changing the notebook’s title, and adding metadata tags from resources such as the Disease Ontology^30^, the Drug Ontology^31^, and the Uber-anatomy Ontology^32^. Once this process is complete, a notebook generation job is launched by the BioJupies notebook generator server. The BioJupies notebook generator creates and executes a Jupyter Notebook file with the uploaded, or fetched, dataset and selected plug-ins. When the process is completed, the generated notebook is uploaded to a cloud storage bucket and is made available through a persistent URL (Supplementary Video S3). If the user approves, the persistent URL is made publicly and automatically available on the Datasets2Tools repository^33^. A schematic representation of the entire BioJupies workflow shows how the various components of BioJupies come together (Fig. 1).

**Fig.1.**
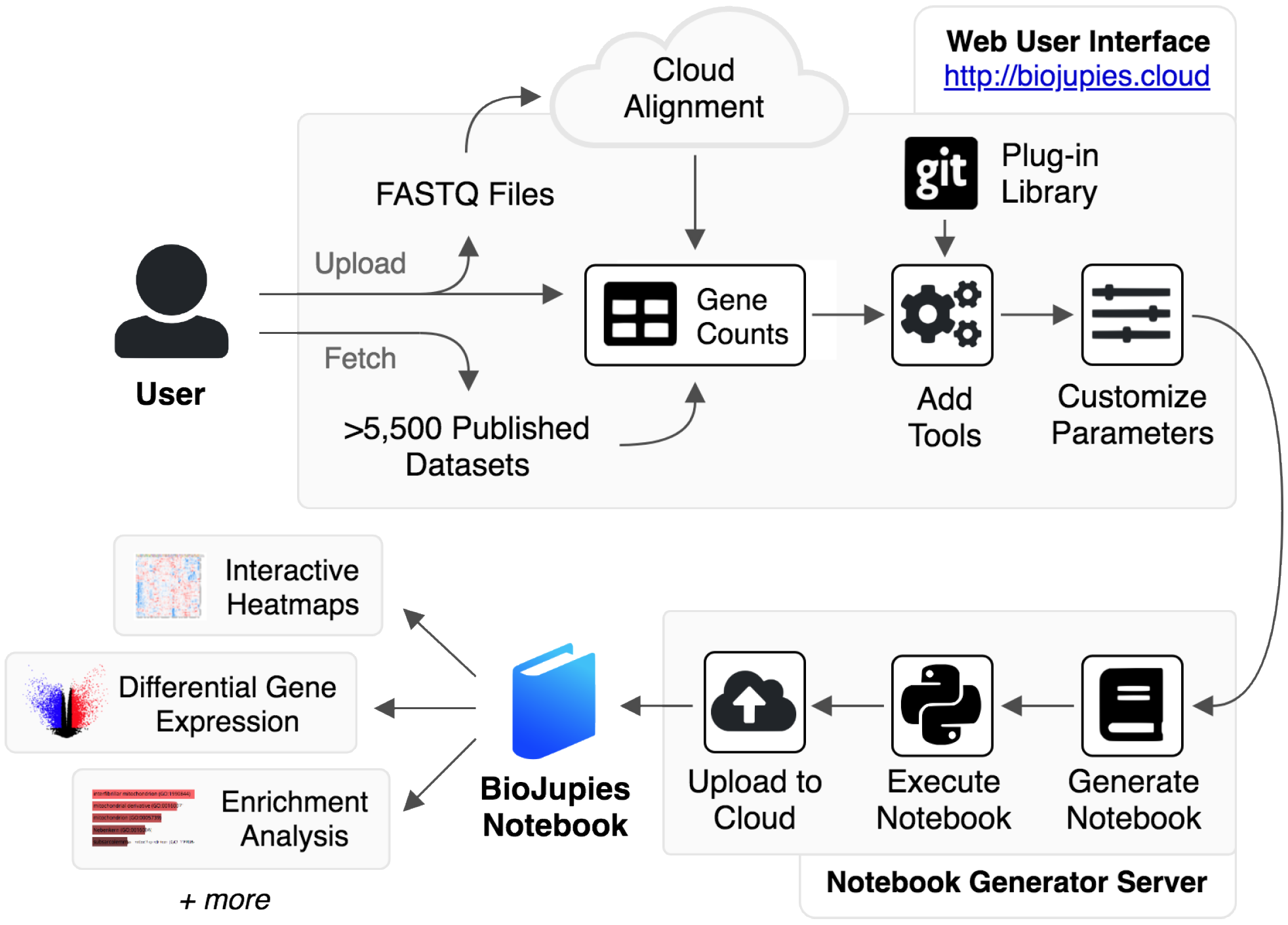
Schematic illustration of the BioJupies notebook generation workflow. The user starts by uploading RNA-seq data to the BioJupies website (http://biojupies.cloud), or by selecting from thousands of publicly available datasets. If raw FASTQ files are provided, expression levels for each gene are quantified using a cloud-based alignment pipeline. The user subsequently selects the tools and parameters to apply to analyze the data. Finally, a server generates a Jupyter Notebook with the desired settings and returns a report to the user through a persistent URL.

### Uploading RNA-seq Data

BioJupies enables the generation of Jupyter Notebooks from RNA-seq data in both raw and processed forms. In case of processed RNA-seq data, the user uploads numeric gene counts in a tabular format. This can be an Excel spreadsheet or a comma-separated text file. In addition, metadata that describes the samples can be uploaded in separate Excel spreadsheets, or as comma-separated text files, following the provided examples. In the case of raw RNA-seq data, the user is provided with a user interface that enables them to upload FASTQ files directly to an AWS S3 cloud storage bucket through an HTML form (Supplementary Video S1). The user is required to specify the organism, and whether the RNA-seq data was generated using single-end or paired-end sequencing. Once this information is collected, gene expression levels for each gene are quantified by launching parallel alignment jobs in the cloud using the ARCHS4 pipeline. The pipeline employs the kallisto aligner for the alignment step^29^. We have benchmarked the kallisto aligner with other aligners and found it to produce comparable read quality at a significant lower cost^2^. Once the alignment process is complete, which may take up to 10 minutes, sample counts are merged to generate a gene count matrix. The gene count matrix is subsequently combined with the uploaded metadata. From that point on, the user follows the same steps to generate notebooks as with fetched data, or processed uploaded data (gene counts matrix), by selecting the computational tools they wish to employ (Supplementary Video S2).

### Analyzing Published Datasets

In addition to allowing analysis of user-submitted data, BioJupies provides direct access to data from >5,500 published GEO studies. These studies contain >250,000 processed samples from human or mouse, profiled across hundreds of different tissue types, cell lines and biological conditions. The BioJupies website enables users to fetch the data into BioJupies from these studies using a search engine that supports text-based queries. The search engine has filters, for example, by specifying a range of number of samples per study, filtering by organism, and filtering by a publication date range. The datasets available for fetching are updated regularly as more studies are published in GEO.

### The BioJupies Plug-in Library

All the source code that is used to analyze the RNA-seq data and generate the plots is available on GitHub at a public repository: https://github.com/MaayanLab/biojupies-plugins. Currently, the BioJupies plug-in library consists of 14 computational plug-ins subdivided into four categories: exploratory data analysis, differential expression analysis, enrichment analysis, and small molecule query (Table 1). These plug-ins analyze the data and display interactive visualizations such as three-dimensional scatter plots, bar charts, and clustered heatmaps with enrichment analysis features (Supplementary Video S3). The plug-in’s modular design of BioJupies provides the mechanism needed for other bioinformatics tool developers to integrate their tools with BioJupies. Suggestions for plug-ins can be submitted to the BioJupies web-site or through GitHub.

### The BioJupies Chrome Extension

BioJupies is also provided as a Chrome extension. The Chrome extension is freely available from the Chrome Web Store at: https://chrome.google.com/webstore/detail/biojupies-generator/picalhhlpcjhonibabfigihelpmpadel?hl=en-US. The Chrome extension detects BioJupies available processed datasets returned from GEO search pages. The BioJupies Chrome extension embeds buttons near each entry from returned results of GEO studies that were aligned by us. Clicking on the embedded button, a popup window is triggered to provide an interactive interface to capture the settings to generate the Jupyter Notebook directly from the selected GEO dataset. The BioJupies Chrome extension entry forms follow the same content as in the BioJupies website. The BioJupies Chrome extension launches a notebook generation job and displays the permanent link to the user once the process is completed (Supplementary Video S4).

## Discussion

The automated generation of Jupyter Notebooks for RNA-seq data analysis lowers the point-of-entry for researchers with no programming background. With BioJupies users can rapidly extract knowledge from their own RNA-seq data, or from already published studies. Furthermore, the notebooks generated can be easily shared with collaborators, rerun and customized using Docker containers to further refine the analysis. This makes it attractive to both experimentalists and programmers. The pipeline is cost effective and scalable, so it can accommodate a large user base. However, if hundreds of users utilize the service daily, the costs can start mounting to levels that are not sustainable by us. In such case, technologies that transfer some of the cloud computing costs to the users may be needed.

It may appear that the RNA-seq Jupyter Notebooks make the RNA-seq data, and the tools applied for the analysis, more findable, accessible, interoperable and reusable (FAIR)^34^. However, currently, the notebooks and the datasets analyzed by BioJupies fail many of the FAIR guidelines and metrics. For example, the notebook is not citable, uploaded RNA-seq data is not indexed in an established repository, metadata describing the samples and experimental conditions are not required, and unique identifiers are not enforced. Future work will enhance the process to better comply the BioJupies generated notebooks with the FAIR principles. One way to increase FAIRness is to encode the BioJupies pipelines in standard workflow languages such as Common Workflow Language (CWL)^35^ or Workflow Description Language (WDL)^36^. This approach may facilitate the more rapid adoption of other data analysis pipelines that can be supported by the BioJupies framework. Although BioJupies was developed to create notebooks for RNA-seq analysis, pipelines to process other data types should be possible to implement. This can be achieved through the plug-in architecture and coded workflows. One feature that is currently missing from BioJupies is the launch of live notebooks in the cloud. To achieve this, Kubernetes^37^ can be utilized to directly deploy BioJupies in the cloud. Hence, BioJupies can be extended in many ways. While there are many ways to extend BioJupies, at its present form, the BioJupies application can facilitate rapid and in-depth data analysis for many investigators.

## Acknowledgements

This work is supported by NIH grants U54-HL127624 (LINCS-DCIC), U24-CA224260 (IDG-KMC), and OT3-OD025467 (NIH Data Commons).

## Supplementary Videos

**Supplementary Video 1. Uploading and aligning raw RNA-seq data using BioJupies.** https://www.youtube.com/watch?v=mZR02DCv8RE

**Supplementary Video 2. Generating a notebook using the BioJupies website.** https://www.youtube.com/watch?v=FpeGHhTm9Hg

**Supplementary Video 3. Exploring the contents of a BioJupies notebook.**
https://www.youtube.com/watch?v=dApCOur2Bbo

**Supplementary Video 4. Generating a notebook using the BioJupies Chrome extension.** https://www.youtube.com/watch?v=164zkRKsC18

## References

1. Wang, Z., Gerstein, M. & Snyder, M. RNA-Seq: a revolutionary tool for transcriptomics. Nat. Rev. Genet. 10, 57–63 (2009).

2. Lachmann, A. et al. Massive mining of publicly available RNA-seq data from human and mouse. Nat. Commun. 9, 1366 (2018).

3. Kluyver, T. et al. Jupyter Notebooks – a publishing format for reproducible computational workflows. in Positioning and Power in Academic Publishing: Players, Agents and Agendas (eds. by Loizides, F. & Scmidt, B.) 87–90 (IOS Press, 2016).

4. Knuth, D. E. Literate Programming. Comput. J. 27, 97–111 (1984).

5. RStudio Team. RStudio: Integrated Development Environment for R. (RStudio, Inc., 2015).

6. Xie, Y. et al. knitr: A General-Purpose Package for Dynamic Report Generation in R. (2018).

7. https://beta.observablehq.com/. Observable. Available at: https://beta.observablehq.com/. (Accessed: 25th May 2018)

8. https://zeppelin.apache.org/.Zeppelin. Available at: https://zeppelin.apache.org/. (Accessed: 21st May 2018)

9. Wang, Z. & Ma’ayan, A. An open RNA-Seq data analysis pipeline tutorial with an example of reprocessing data from a recent Zika virus study. F1000Research 5, 1574 (2016).

10. Dumontier, M. & Kuhn, T. Data Science & nbsp;–Methods, infrastructure, and applications. Data Sci. 1, 1–5 (2017).

11. Qu, K. et al. Integrative genomic analysis by interoperation of bioinformatics tools in GenomeSpace. Nat. Methods 13, 245–247 (2016).

12. Afgan, E. et al. The Galaxy platform for accessible, reproducible and collaborative biomedical analyses: 2018 update. Nucleic Acids Res. (2018). doi:10.1093/nar/gky379

13. Reich, M. et al. GenePattern 2.0. Nat. Genet. 38, 500–501 (2006).

14. Grüning, B. A. et al. Jupyter and Galaxy: Easing entry barriers into complex data analyses for biomedical researchers. PLOS Comput. Biol. 13, e1005425 (2017).

15. Reich, M. et al. The GenePattern Notebook Environment. Cell Syst. 5, 149–151.e1 (2017).

16. Chang, W. et al. shiny: Web Application Framework for R. (2018).

17. Ge, S. X. iDEP: An integrated web application for differential expression and pathway analysis. bioRxiv 148411 (2017). doi:10.1101/148411

18. Nelson, J. W., Sklenar, J., Barnes, A. P. & Minnier, J. The START App: a web-based RNAseq analysis and visualization resource. Bioinformatics 33, 447–449 (2017).

19. Younesy, H., Möller, T., Lorincz, M. C., Karimi, M. M. & Jones, S. J. VisRseq: R-based visual framework for analysis of sequencing data. BMC Bioinformatics 16, S2 (2015).

20. Gardeux, V., David, F. P. A., Shajkofci, A., Schwalie, P. C. & Deplancke, B. ASAP: a web-based platform for the analysis and interactive visualization of single-cell RNA-seq data. Bioinformatics 33, 3123–3125 (2017).

21. Li, Y. & Andrade, J. DEApp: an interactive web interface for differential expression analysis of next generation sequence data. Source Code Biol. Med. 12, 2 (2017).

22. Monier, B., McDermaid, A., Zhao, J., Fennell, A. & Ma, Q. IRIS-DGE: An integrated RNA-seq data analysis and interpretation system for differential gene expression. bioRxiv 283341 (2018)

23. Edgar, R., Domrachev, M. & Lash, A. E. Gene Expression Omnibus: NCBI gene expression and hybridization array data repository. Nucleic Acids Res. 30, 207–210 (2002).

24. Merkel, D. Docker: Lightweight Linux Containers for Consistent Development and Deployment. Linux J 2014, (2014).

25. McKinney, W. Data Structures for Statistical Computing in Python. in Proceedings of the 9th Python in Science Conference (eds. van der Walt, S. & Millman, J.) 51–56 (2010).

26. Ritchie, M. E. et al. limma powers differential expression analyses for RNA-sequencing and microarray studies. Nucleic Acids Res. 43, e47–e47 (2015).

27. Clark, N. R. et al. The characteristic direction: a geometrical approach to identify differentially expressed genes. BMC Bioinformatics 15, 79 (2014).

28. The HDF5® Library & File Format. The HDF5® Library & File Format. The HDF Group

29. Bray, N. L., Pimentel, H., Melsted, P. & Pachter, L. Near-optimal probabilistic RNA-seq quantification. Nat. Biotechnol. 34, 525–527 (2016).

30. Kibbe, W. A. et al. Disease Ontology 2015 update: an expanded and updated database of human diseases for linking biomedical knowledge through disease data. Nucleic Acids Res. 43, D1071–1078 (2015).

31. Hanna, J., Joseph, E., Brochhausen, M. & Hogan, W. R. Building a drug ontology based on RxNorm and other sources. J. Biomed. Semant. 4, 44 (2013).

32. Mungall, C. J., Torniai, C., Gkoutos, G. V., Lewis, S. E. & Haendel, M. A. Uberon, an integrative multi-species anatomy ontology. Genome Biol. 13, R5 (2012).

33. Torre, D. et al. Datasets2Tools, repository and search engine for bioinformatics datasets, tools and canned analyses. Sci. Data 5, 180023 (2018).

34. Wilkinson, M. D. et al. The FAIR Guiding Principles for scientific data management and stewardship. Scientific Data (2016). doi:10.1038/sdata.2016.18

35. Amstutz, P. et al. Common Workflow Language, v1.0. (2016).

36. https://software.broadinstitute.org/wdl/. WDL | Home. Available at: https://software.broadinstitute.org/wdl/. (Accessed: 11th June 2018)

37. Hightower, K., Burns, B. & Beda, J. Kubernetes: Up and Running Dive into the Future of Infrastructure. (O’Reilly Media, Inc., 2017).

38. Pedregosa, F. et al. Scikit-learn: Machine Learning in Python. J. Mach. Learn. Res. 12, 2825–2830 (2011).

39. https://plot.ly. Modern Visualization for the Data Era. undefined Available at: https://plot.ly. (Accessed: 21st May 2018)

40. Fernandez, N. F. et al. Clustergrammer, a web-based heatmap visualization and analysis tool for high-dimensional biological data. Sci. Data 4, 170151 (2017).

41. Kuleshov, M. V. et al. Enrichr: a comprehensive gene set enrichment analysis web server 2016 update. Nucleic Acids Res. 44, W90–97 (2016).

42. Ashburner, M. et al. Gene ontology: tool for the unification of biology. The Gene Ontology Consortium. Nat. Genet. 25, 25–29 (2000).

43. Kanehisa, M., Furumichi, M., Tanabe, M., Sato, Y. & Morishima, K. KEGG: new perspectives on genomes, pathways, diseases and drugs. Nucleic Acids Res. 45, D353–D361 (2017).

44. Slenter, D. N. et al. WikiPathways: a multifaceted pathway database bridging metabolomics to other omics research. Nucleic Acids Res. 46, D661–D667 (2018).

45. Croft, D. et al. The Reactome pathway knowledgebase. Nucleic Acids Res. 42, D472–477 (2014).

46. Lachmann, A. et al. ChEA: transcription factor regulation inferred from integrating genome-wide ChIP-X experiments. Bioinforma. Oxf. Engl. 26, 2438–2444 (2010).

47. Lachmann, A. & Ma’ayan, A. KEA: kinase enrichment analysis. Bioinforma. Oxf. Engl. 25, 684–686 (2009).

48. Chou, C.-H. et al. miRTarBase update 2018: a resource for experimentally validated microRNA-target interactions. Nucleic Acids Res. 46, D296–D302 (2018).

49. Agarwal, V., Bell, G. W., Nam, J.-W. & Bartel, D. P. Predicting effective microRNA target sites in mammalian mRNAs. eLife 4, (2015).

50. Duan, Q. et al. L1000CDS^2^: LINCS L1000 characteristic direction signatures search engine. Npj Syst. Biol. Appl. 2, 16015 (2016).

51. Wang, Z., Lachmann, A., Keenan, A. B. & Ma’ayan, A. L1000FWD: Fireworks visualization of drug-induced transcriptomic signatures. Bioinforma. Oxf. Engl. (2018).

